# Opioids trigger breast cancer metastasis through E-Cadherin downregulation and STAT3 activation promoting epithelial-mesenchymal transition

**DOI:** 10.1101/443663

**Authors:** Sabrina Tripolt, Vanessa M. Knab, Heidi A. Neubauer, Dominik P. Elmer, Fritz Aberger, Richard Moriggl, Daniela A. Fux

## Abstract

The opioid crisis of pain medication bears risks from addiction to cancer progression, but little experimental facts exist. Expression of δ-opioid receptors (DORs) correlates with poor prognosis for breast cancer (BCa) patients, but mechanism and genetic/pharmacologic proof of key changes in opioid-triggered cancer biology are lacking. We show that oncogenic STAT3 signaling and E-Cadherin downregulation are triggered by opioid-ligated DORs, promoting metastasis. Human and murine transplanted BCa cells (MDA-MB-231, 4T1) displayed enhanced metastasis upon opioid-induced DOR stimulation, and DOR-antagonist blocked metastasis. Opioid-exposed BCa cells showed enhanced migration, STAT3 activation, down-regulation of E-Cadherin and expression of epithelial-mesenchymal transition (EMT) markers. STAT3 knockdown or upstream inhibition through the JAK1/2 kinase inhibitor ruxolitinib prevented opioid-induced BCa cell metastasis and migration. We conclude that opioids trigger metastasis through oncogenic JAK1/2-STAT3 signaling.

## Introduction

Opioids are potent analgesic drugs that are indispensable in cancer therapy as their use alleviates pain during and after tumor resection surgery. Moreover, opioids are applied to relieve cancer-related pain resulting from the tumor itself pressing on different organs. Opioids are also administered to manage pain arising from chemo-or radiotherapy (1, 2). However, opioids are also widely misused and often given to risk group patients of older age with chronic disease/pain who may be more susceptible to developing invasive cancers. The use of opioids in cancer patients is a controversial debate. Epidemiologic studies revealed that administering opioids after surgical tumor removal enhanced the risk of cancer recurrence or enhanced metastasis in BCa patients undergoing mastectomy and axillar clearance (3–5). In addition, opioid treatment of patients with advanced malignant tumors is associated with worse overall survival (6–9). Enhanced metastasis was also reported in opioid-treated rodents with mammary carcinomas (10, 11), but the cellular and molecular mechanisms underlying these observations remain unclear.

The analgesic effects of opioids are mediated by μ-, δ- and κ-opioid receptors, which belong to the family of G protein-coupled receptors (GPCRs). Opioid receptors are linked to different intracellular signaling cascades including ERK1/2 and AKT signaling (12), which contribute to various cellular processes including proliferation, differentiation, and survival (13, 14). Opioid receptors are most strongly expressed in neurons of the nociceptive system, but can also be found in heart, immune system, gastrointestinal tract and reproductive system cells (15–17). Recently, high expression of δ-opioid receptors (DOR) was observed in tissue samples from BCa patients, which correlated with tumor progression and poor prognosis (9). Moreover, DOR expression was closely associated with tumor aggressiveness and distant metastasis. As the findings imply a critical involvement of DOR signaling in cancer cell dissemination, we investigated the role of DOR signaling on migration and metastasis of mammary tumor cells. We explored the mechanism behind using mammary BCa cell line transplant systems with human and murine BCa cell lines and pharmacologic as well as genetic evaluation of essential players found to be hyperactivated upon opioid-triggered DOR activation. Our mechanistic insights reveal that DOR activation promotes JAK1/2-STAT3 activation, thereby inducing EMT and accelerating breast cancer metastasis.

## Results

### Expression of δ-opioid receptors and association with solid tumor progression

To obtain a comprehensive view of the role of DOR in BCa disease progression and metastasis, gene expression analyses were performed using BCa patient datasets publicly available from the Oncomine database. As shown in **Fig. 1A**, the expression of DOR (*OPRD1*) mRNA was higher in Grade 2 than in Grade 1 mammary malignancies. Moreover, Stage III BCa exhibited higher DOR expression than Stage II tumors. Notably, expression of DOR mRNA correlated with enhanced disease progression in BCa metastasis (**Fig. 1A**). Interestingly, these observations are not restricted to mammary malignancies. Oncomine data evaluation also revealed significant associations of increased DOR expression and cancer recurrence/metastasis/survival in squamous cell carcinomas of the tongue or head and neck, malignant melanoma or ovarian cancer (**Fig. S1A-D**). Increased DOR expression was also found in lung adenocarcinoma and cancers of testis, renal, uterus cells types and melanoma versus healthy control tissues (**Fig. S1E-J**).

**Figure 1.**
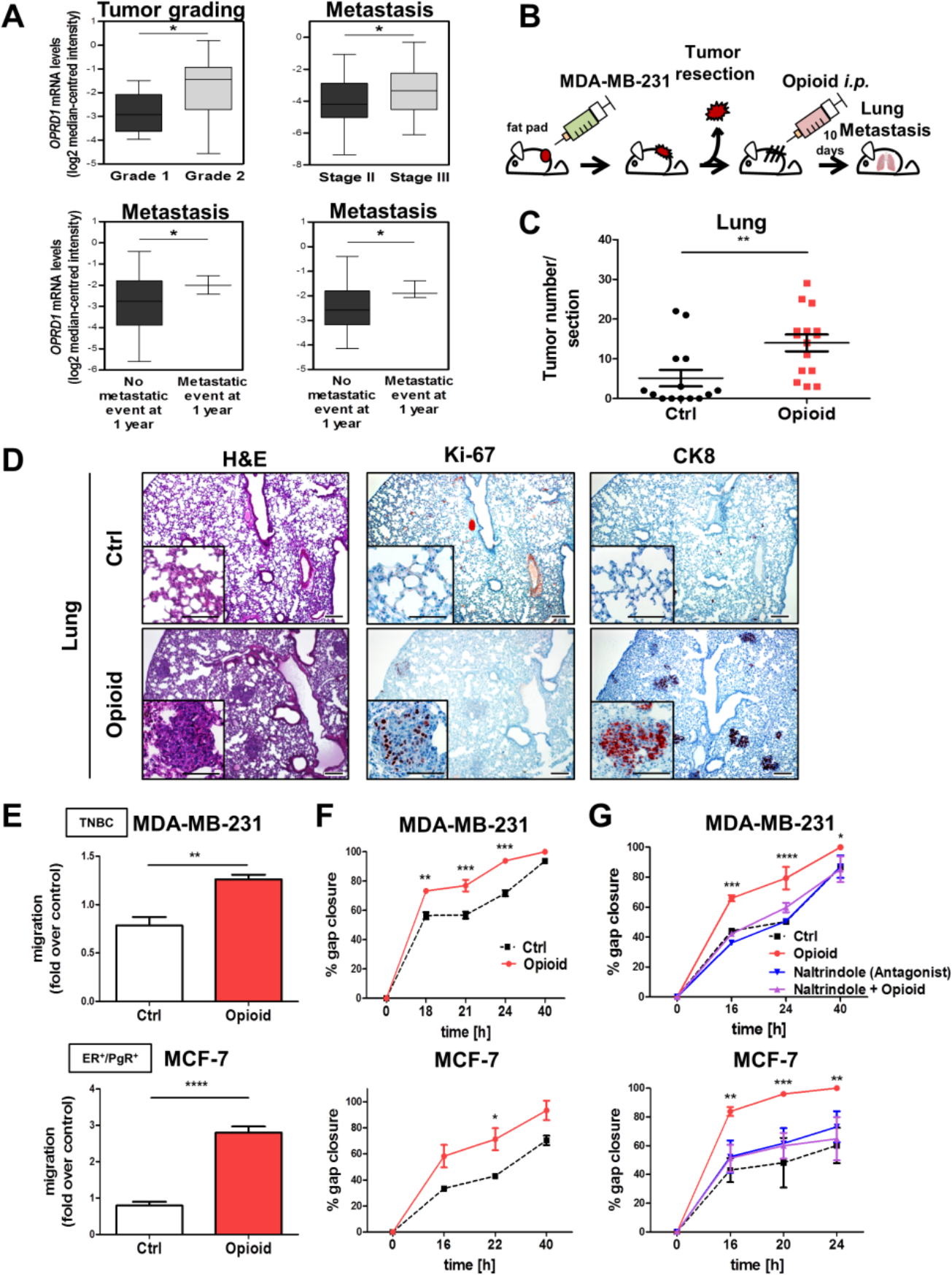
Role of δ-opioid receptors in breast cancer metastasis. (**A**) Relationship between *OPRD1* mRNA and mammary tumor progression were examined using the Oncomine database. **(A; upper left)** Comparison of *OPRD1* expression in BCa patients Grade 1 and Grade 2. **(A; upper right)** Comparison of *OPRD1* expression in BCa patients Stage II and Stage III. (**A; lower panels**) Comparison of *OPRD1* expression in BCa patients having no metastatic events at 1 year and metastatic events at 1 year. (Datasets: Desmedt Breast or Minn Breast 2; ^*^P<0.05, Student’s t-test). **(B)** Experimental scheme. **(C)** Quantification of metastasis in the lungs of opioid- and saline-treated Rag2^-/-^γc^-/-^ mice. Graphs represent quantification of CK8-positive tumors per section as mean ± SD (n=14 for PBS-treated mice and n=15 for opioid-treated mice; ^**^P<0.01; Student’s t-test). **(D)** Immunostainings of lungs from control and opioid-treated mice for H&E, Ki-67 and CK8 (magnification: 40x and 100x; scale bars 200 μm and inserts 100 μm). **(E)** Transwell migration assay of MDA-MB-231 (upper panel; ^**^P<0.01) and MCF-7 cells (lower panel; ^****^P<0.0001) after opioid stimulation (1 μM, 16 h incubation). Controls (ctrl) were treated with ddH_2_O. (Bar graphs represent the fold change of migrated cells over the control as mean ± SD in duplicates; n=3 experiments, Student’s t-test). **(F)** Scratch assays of MDA-MB-231 and MCF-7 cells exposed to the opioid (1 μM) or ddH2O (ctrl). Quantification represents gap closure (migration) rates as mean ± SD of the % of the closure of original gap (MDA-MB-231: ^**^P<0.01 versus control; ^***^P<0.001 versus control; MCF-7: ^*^P<0.05 versus control; n=3 experiments, in duplicates, two-way ANOVA). **(G)** Scratch assay of MDA-MB-231 and MCF-7 cells treated with 1 μM opioid in the absence or presence of 10 μM naltrindole (MDA-MB-231: ^*^P<0.05 versus control; ^***^P<0.001 versus control; ^****^P<0.0001 MCF-7: ^**^P<0.01 versus control; ^***^P<0.001 versus control; n=3 experiments, in duplicates, two-way ANOVA).

The striking data on BCa progression and the clinical need to explain BCa progression prompted us to further explore the role of DOR in metastasis using a BCa transplant model. MDA-MB-231 cells were orthotopically implanted into mammary fat pads of immunocompromised *Rag2^-/-^γc^-/-^* mice. After surgical removal of the primary tumor, mice were treated with a classic DOR agonist [D-Ala^2^,D-Leu^5^]-Enkephalin (further indicated as opioid) for ten days, and metastases were examined in the lungs of the mice by immunohistochemistry (**Fig. 1B**). Compared to the vehicle control, lungs of opioid-treated mice showed more clusters of Ki-67 and Cytokeratin 8 (CK8) positive cells (**Fig. 1C, 1D**), identifying them as metastatic BCa cells (18). As an additional immunocompetent model, we performed opioid treatment in Balb/C wildtype mice bearing murine 4T1 BCa cell transplants, which also resulted in more metastatic lung nodules than vehicle controls (**Fig. S2A**). Results from these two transplant models demonstrate that opioid treatment increased lung metastasis of primary BCa cells.

To gain insights into the mechanism of opioid-induced metastasis, migration of DOR-expressing BCa cells was examined. Three human BCa cell line models were used MDA-MB-231, MCF-7 and T47D cells, all of which endogenously express DOR (**Fig. S2B**). Transwell and scratch assays showed that exposure of MDA-MB-231, MCF-7 and T47D BCa cells to opioid enhanced migration (**Fig. 1E, S2C**, **1F** and **S2D**). Opioid-induced migration was abolished by naltrindole, a DOR-selective antagonist (19) (**Fig. 1G and S2E**). Importantly, opioid exposure neither affected proliferation nor cell cycle distribution (**Fig. S2F, S2G**). Altogether, these findings indicate that opioids promote BCa cell migration by DOR stimulation.

### STAT3 is activated by a DOR agonist in breast cancer cells

Migration can be promoted through multiple core cancer pathways, including the JAK-STAT3/5, β-Catenin/WNT, RAS/RAF-ERK1/2 or TGF-β/SMAD2/3 signaling pathways. Whereas incubation of MCF-7, T47D and MDA-MB-231 cells with opioid for different time periods had no significant effect on activation of β-Catenin (ABC), ERK1/2 or SMAD2/3 (**Fig. S3A**), immunoblotting showed that cell exposure to opioid resulted in increased STAT3 activation (pYSTAT3) (**Fig. 2A, S3B**). In contrast STAT5A/B transcription factors also known to promote BCa progression (20), remained unaffected (**Fig. S3C**).

STAT3 can control cell migration through promotion of EMT (21). To test whether opioid treatment can induce EMT, BCa cells were analyzed for classical EMT-related genes controlled by STAT3, in particular the E-Cadherin repressors SNAIL, SLUG and TWIST (22–25). qRT-PCR analysis and immunoblotting revealed that opioid treatment led to an up-regulation of *SNAIL* at both the mRNA and protein level in all tested BCa cells (**Fig. 2A, 2B, S3D and S3E**). Moreover, TWIST protein expression was enhanced in opioid-treated MCF-7 and T47D cells, whereas *SLUG* was up-regulated in MDA-MB-231 cells. FACS analysis yielded a loss of E-Cadherin protein on the surface of all tested BCa cells (**Fig. 2C, S3E)**. Moreover, E-Cadherin mRNA was decreased in MCF-7 and MDA-MB-231 cells. Immunostainings of lungs from opioid-treated mice bearing BCa xenografts further demonstrated that the observed metastases are highly positive for Vimentin, a further EMT marker, but negative for Cleaved Caspase-3 (**Fig. 2D**). Therefore, our data suggest that opioids promote EMT through STAT3 activation, loss of E-Cadherin and gained Vimentin expression.

**Figure 2.**
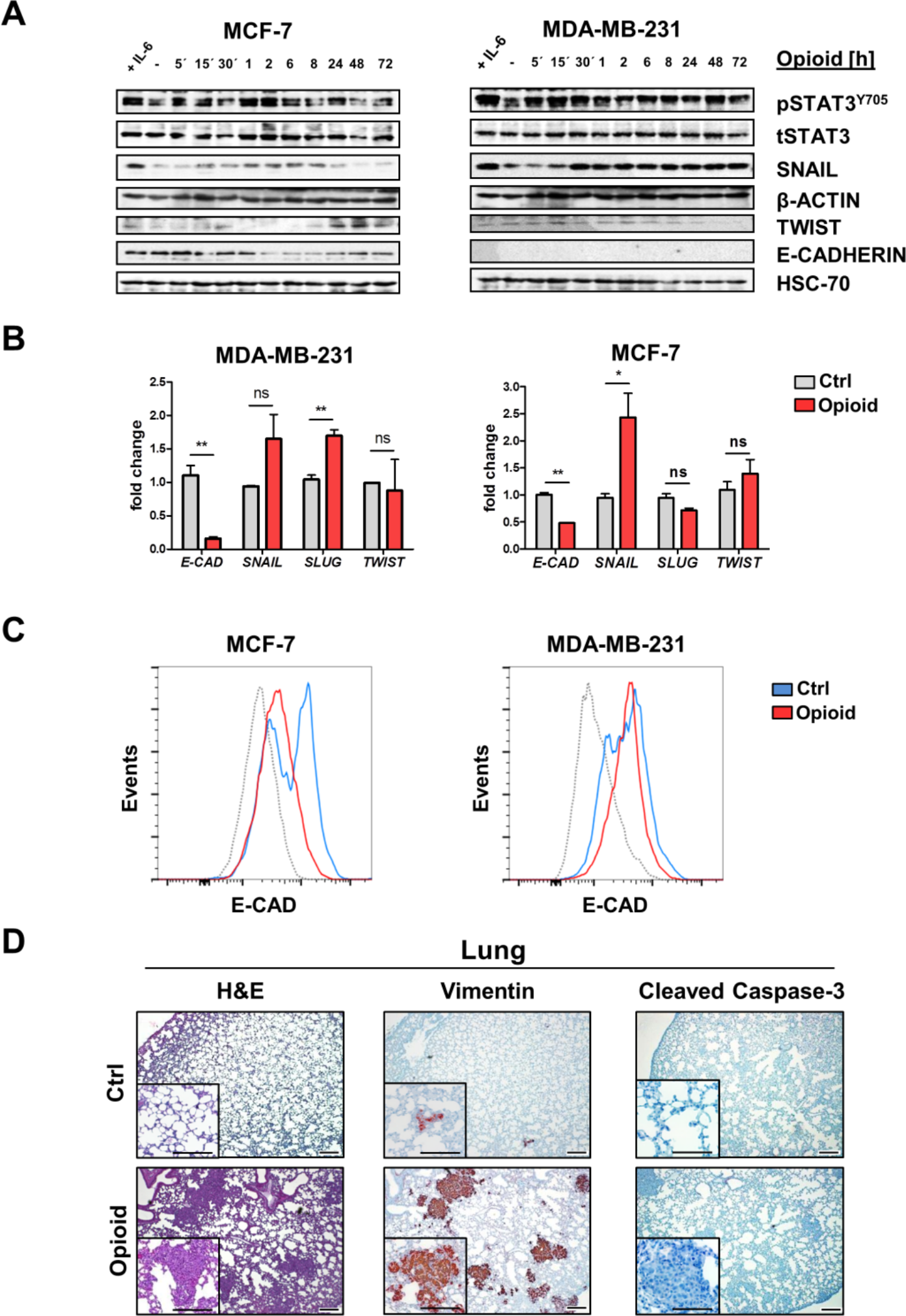
Opioid exposure induces STAT3 tyrosine phosphorylation and expression of individual EMT-related marker in breast cancer cells. **(A)** MDA-MB-231 and MCF-7 cells were treated with 1 μM opioid for 5 min – 72 h and analyzed for pSTAT3^Y705^, tSTAT3, SNAIL, TWIST and E-CADHERIN levels by immunoblotting. β-ACTIN and HSC-70 were used as loading controls. (**B**) MDA-MB-231 and MCF-7 cells were treated with 1 μM opioid for 1 h and examined for mRNA levels of *E-Cadherin*, *SNAIL*, *SLUG* and *TWIST* by real-time qRT-PCR (^*^p<0.05, ^**^p<0.01, n=3 experiments, Student’s t-test). **(C)** FACS histogram plots (blue: ctrl treated, red: opioid-treated 1 μM, 1 h) of MCF-7 and MDA-MB-231 cells stained against E-Cadherin. Dotted curves represent FACS histogram plots for the isotype-matched controls. **(D)** Representative H&E, Cleaved Caspase-3 and Vimentin stained lung sections of mice engrafted with MDA-MB-231 cells after primary tumor removal and post-surgical opioid treatment for ten days (original magnification: 40x and 100x; scale bars 200 μm and inserts 100 μm).

### Role of STAT3 in opioid-mediated cell migration and metastasis

Next, the role of STAT3 activation in opioid-mediated cell migration was examined. First, cell motility was assessed in the presence of the FDA-approved JAK1/2 inhibitor ruxolitinib. Ruxolitinib blocked opioid-induced STAT3 activation (**Fig. 3A**) and prevented opioid-triggered migration (**Fig. 3B, S4A**). Opioid-induced STAT3 activation was also blocked by the DOR antagonist naltrindole (**Fig. 3A**). In a second approach, STAT3 function was blocked by stable RNA knockdown. Infection of BCa cells with *STAT3-shRNA* reduced STAT3 expression by more than 70% at both the mRNA and protein level (**Fig. S4B, S4C**). Importantly, *STAT3* knockdown did not affect cell proliferation, cell cycle distribution or basal migration activity (**Fig. S4D-F**), but notably it prevented increased migration of BCa cells in response to opioid exposure (**Fig. 3C, S4G**).

**Figure 3.**
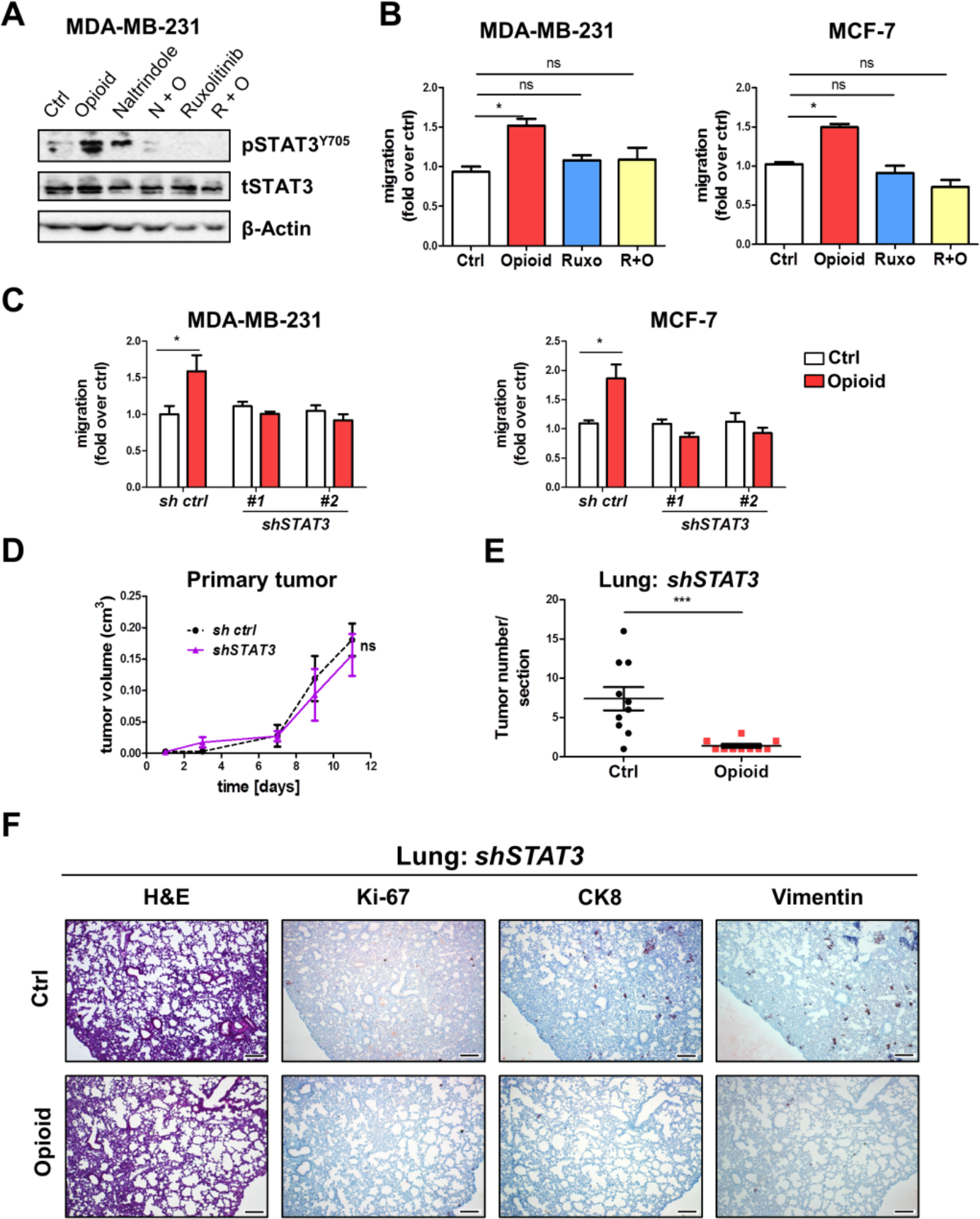
Inhibition of STAT3 attenuates opioid-triggered breast cancer cell migration and metastasis. **(A)** MDA-MB-231 cells were treated with 1 μM opioid in the absence or presence of naltrindole (10 μM) or ruxolitinib (3.3 nM), and analyzed for pSTAT3^Y705^ and tSTAT3 by immunoblotting. β-ACTIN was used as loading control. **(B)** Transwell assay. MDA-MB-231 and MCF-7 were exposed to opioid (1 μM) alone or together with ruxolitinib (3.3 nM). After 16 h, cells that migrated through the membrane pores were quantified (^*^P<0.05 opioid versus vehicle control (DMSO); n=3 experiments, one-way ANOVA). **(C)** MDA-MB-231 and MCF-7 cells were infected with unrelated *control-shRNA* (*sh ctrl*), *STAT3-shRNA* #1, or *STAT3-shRNA* #2, and examined for migration activity after opioid treatment by transwell assay. Controls were treated with ddH_2_O (^*^P<0.05 opioid versus control; n=3 experiments, two-way ANOVA). **(D)** Volumes of primary tumors formed by MDA-MB-231 cells infected with *sh ctrl* or *STAT3-shRNA* (*shSTAT3*) over time (not statistically significant (ns), two-way ANOVA). **(E)** Quantification of tumor metastasis in the lungs of opioid-treated *Rag2^-/-^γc^-/-^* mice engrafted with *STAT3-shRNA* infected MDA-MB-231 cells. Graphs depict CK8-positive tumor spots per section as mean ± SD (n=10 for PBS-treated mice and n=10 for opioid-treated mice; ^***^P<0.001, Student’s t-test). **(F)** Representative images of H&E, Ki-67, CK8 and Vimentin immunostainings of lungs from mice engrafted with STAT3-silenced MDA-MB-231 cells after primary tumor removal and post-surgical opioid treatment for ten days. Control mice were treated with PBS (vehicle).

To also assess the role of STAT3 in opioid-driven metastasis, MDA-MB-231 cells stably transduced with *STAT3-shRNA* or *control-shRNA* were orthotopically injected into female *Rag2^-^ /-γc^-/-^* mice. Primary tumors formed by STAT3-silenced cells were similar in growth kinetics and volume compared to MDA-MB-231 control-shRNA cells (**Fig. 3D**). Following primary tumor resection, mice were treated with opioid or vehicle and lung metastasis was examined. Strikingly and in line with *in vitro* results, animals injected with *STAT3-shRNA* infected cells had significantly less CK8-positive metastatic lesions in the lungs after opioid treatment than vehicle-treated controls (**Fig. 3E, 3F**). Lungs from opioid-treated mice injected with STAT3-silenced cells also displayed significantly less Ki-67 and Vimentin staining than saline controls (**Fig. 3F**). Taken together, our results show that opioid-triggered migration and metastasis of BCa cells requires JAK1/2-STAT3 signaling.

## Discussion

The use of opioids in cancer patients is currently under scrutiny as evidence is emerging of adverse effects on tumor progression. Even worse, people take opioids as pain killers alleviating chronic disease symptoms, which led to an ongoing debate for opioid safety questioning if Western societies are in an opioid crisis. In this study we warn from the detrimental biological function of opioids in cancer progression as evidenced by their capacity to boost migration and metastasis through the induction of an EMT program. We data mined public datasets that display DOR expression to be upregulated in many solid cancers, focusing further on advanced and invasive BCa, where DOR expression levels correlated with accelerated metastatic disease. Indeed, using *in vitro* techniques and different BCa transplant models in immunosuppressed and immunocompetent mice, we demonstrate that DOR activity accelerates BCa migration and metastasis. Furthermore, our pharmacologic and genetic interference studies pinpoint an essential role of JAK1/2-STAT3 activation in opioid-induced EMT processes through E-Cadherin expression blockade.

Metastasis is a multistep process which includes dissociation of single tumor cells from the primary tumor, invasion into the adjacent tissue, immune cell escape, transendothelial migration to enter and leave the circulation system, and subsequent proliferation to colonize within distinct organs (26). Therefore, opioids may promote BCa metastasis by various mechanisms. Our tissue culture experiments revealed that DOR stimulation enhances BCa cell migration, suggesting that opioid-promoted metastasis originates from enhanced cancer cell motility. DOR-stimulated cell migration has been previously observed for non-tumor cells such as epithelial cells, fibroblasts and keratinocytes (27–29). Interestingly, we found that opioid exposure failed to induce classical migration pathways such as MAPK/ERK, TGF-β/SMAD2/3 or WNT/β-Catenin in BCa cells, but activated migratory STAT3 signaling (30, 31) to increase the metastatic potential of tumor cells by inducing EMT (21). STAT3 activation without any opioid effect on WNT/β-Catenin was surprising as activation of β-Catenin/WNT signaling is considered an essential driver of cell motility and synergizes in epithelial carcinomas with oncogenic STAT3 activation (32). Our observation thus implies that DORs activate STAT3 signaling in human BCa cells by a mechanism independent of WNT signaling, but dependent on JAK1/2 function. DOR-mediated STAT3 activation was blocked by ruxolitinib a dual-specific JAK1/2 blocker. JAK1/2 kinases are classically associated with cytokine receptors and activated after cytokine-induced stimulation. However, also activation of JAK1/2 by GPCRs was found to be facilitated by direct interaction with the α subunit of G_q_ proteins (33). As DORs are coupled to G_q_ proteins (34), G_αq_ is thus likely to contribute to JAK1/2 and thus STAT3 activation in opioid-exposed BCa cells. However, it was surprising that STAT5 remained unaffected by the opioid, especially as STAT5A and STAT5B are downstream targets of DOR in neuronal cells suggesting that GPCR-mediated STAT3 docking is unique (35, 36). We and others have shown that STAT3 is more highly expressed in BCa cells than STAT5 (37). Thus, it is likely that oncogenic STAT3 action by DOR ligation dominates BCa cell progression in a vicious cycle through EMT induction.

Previous studies revealed that β-endorphin and other selective DOR agonists inhibit thymic and splenic T cell proliferation and cytokine production (38–40), suggesting that DOR signaling could promote metastasis by suppressing immune function. However, BCa metastasis was enhanced in both immunocompetent and immunodeficient mouse models, implying that the immunosuppressive effect of opioids is not essential.

The observed lung metastases in opioid-treated xenografts were all found to be Vimentin positive. Vimentin has a critical role in metastasis by stabilizing mature invadopodia, which is a prerequisite for invasive spread of cancer cells (41). Vimentin is expressed in MDA-MB-231 cells (42, 43), but we found even more intense Vimentin expression in MDA-MB-231 cells after opioid treatment in transplanted tumors. Enhanced Vimentin expression in MDA-MB-231 cells after opioid exposure could be a downstream effect of increased STAT3 activation, as STAT3 can bind to and activate the Vimentin promoter (43). Opioid exposure also resulted in an up-regulation of the EMT marker SNAIL and downregulation of E-Cadherin in BCa cells. Interestingly, opioid treatment led to cell-specific expression of SLUG and TWIST, which suggests that here SNAIL may be the key EMT regulator. SNAIL has been shown to enhance cell movement via RhoB up-regulation, which alters focal adhesion dynamics (44). SLUG stimulates cell motility by remodeling the ACTIN cytoskeleton (45), whereas TWIST was found to activate Focal Adhesion Kinase, which controls migration processes via assembly/disassembly of cell adhesion (46). The loss of E-Cadherin expression is striking and could be a combined consequence of upregulated E-Cadherin repressor SNAIL and STAT3 activation. In contrast, the loss of E-Cadherin from the surface of BCa cells exposed to the opioid for 1 hour suggests a rapid internalization of the transmembrane glycoprotein after DOR stimulation. E-Cadherin is internalized via Clathrin-coated pits (47), which are also used for opioid-induced endocytosis of DORs (48). As stimulated DORs are maximally internalized within 1 h (49), a co-internalization via Clathrin-coated pits might account for the observed loss of E-Cadherin in opioid-exposed BCa cells. Future studies need to broaden our findings to test them on other solid cancers.

Our meta-analysis of gene expression datasets identified that DOR expression significantly correlates with progression and metastasis of human BCa patients. Previous studies revealed that signaling of DOR strongly differs from signaling of μ- and κ-opioid receptors (MOR, KOR). This results in differences in the regulation of nociception and emotional responses (50), but also proliferation, differentiation and survival of cells outside the central nervous system (28, 51, 52). Although it is not clear what causes unique DOR functions, alternative G protein coupling may be relevant. In contrast to MOR and KOR, DOR also activates G_q/11_ proteins (34) and these were found to regulate cancer cell metastasis (53).

Together, our findings demonstrate that opioids promote BCa metastasis by DOR activation. As several clinically relevant opioids such as morphine, methadone, and fentanyl analogues may also trigger DOR signaling (54, 55), application of the potent analgesics in cancer patients or aged patients with enhanced cancer risk has to be considered with more care. We do not propose STAT3 or JAK1/2 inhibition as promising therapy to be combined with opioids, as this could result in severe immunosuppression and a change in stromal-tumor cell signaling (56). Moreover, drugs like ruxolitinib are known to change cancer cell metabolism and may increase the risk of BCa (56). Mechanistic studies of opioid impact on cancer biology reprogramming will be instrumental to circumvent progressive cancers in older patients addicted or misusing opioids also based on prescription by many doctors. In conclusion, we need to redefine strategies to generate for example opioids in cancer-related pain that do not down-regulate E-Cadherin, activate SNAIL or JAK1/2-STAT3 to have safer pain medication that will not promote malignant carcinoma progression.

## Materials and Methods

### Cell culture

BCa cell lines MDA-MB-231, MCF-7, T47D, 4T1 and human glioblastoma cells LN299 were purchased from the American Type Culture Collection (ATCC; VA, USA). LN299 and human BCa cell lines were maintained in Dulbecco’s modified Eagle’s medium (DMEM; Sigma Aldrich, MO, USA) supplemented with 10% fetal bovine serum (FBS; PAA Laboratories Inc., AUT) and 100 U/ml Penicillin-Streptomycin (PAA). 4T1 cells were cultured in RPMI-1640 medium supplemented with 10% FCS and 100 U/ml Penicillin-Streptomycin. Cells were maintained at 37°C with 5% CO_2_.

### RNA interference and lentiviral transduction

Cells with stable *STAT3* knockdown were generated by using lentiviral RNA interference as described in *Kasper et al.* using Metafectene pro (Biontex, GER) transfection reagent. Viral particles were produced in transfected 293FT cells for each of the following short hairpin RNA (shRNA) constructs (Mission TRC shRNA library, Sigma Aldrich): control shRNA (SHC002), shRNA *STAT3* (shSTAT3) #1 (TRCN0000071456) and #2 (TRCN0000020843). Lentiviral transduction of MCF-7, MDA-MB-231, and T47D cells with control and shSTAT3 constructs was performed as previously described (57). Two days after transduction, BCa cells were selected with puromycin and further analyzed for knockdown efficiency by Western blot and qPCR.

### Cell extracts and immunoblotting

Immunoblotting was performed as previously described (58). Briefly, cells were treated with 1 μM [D-Ala^2^, D-Leu^5^]-Enkephalin acetate salt (DADLE; Bachem, CH; in text indicated as opioid) as indicated and lysed with ice-cold RIPA buffer (10 mM Tris-Cl (pH 8.0), 1 mM EDTA, 0.5 mM EGTA, 1% Triton X-100, 0.1% sodium deoxycholate, 0.1% SDS, 140 mM NaCl). Total cell lysates (30 μg) were subjected to 7% SDS polyacrylamide gel electrophoresis and blotted on nitrocellulose membrane (GE Healthcare, GBR). Membranes were blocked with 5% BSA in TBS/Tween-20 (0.1%) and probed with antibodies against active-β-Catenin (ABC) (Cell Signaling Technology (CST), MA, USA; 95625), E-Cadherin (14472), GAPDH (2118), phospho-p42/p44 (Thr202/Tyr204) (9101), total SMAD2/3 (8685), phospho-Smad2 (Ser465/467)/Smad3 (Ser423/425) (8828), Snail (3879), phospho-STAT3 (Tyr705) (9131), total p42/p44 (9102), total STAT3 (12640), total STAT5 (9363), phospho-STAT5 (Tyr694) (Becton Dickinson (BD), NJ, USA; 611964), DOR-1 (Santa Cruz Biotechnology (SCB), TX, USA; 9111), HSC-70 (7298), TWIST (81417) or β-ACTIN (69879). After washing and membrane incubation with respective secondary antibodies, proteins were detected by chemiluminescence using LumiGLO Reagent (CST) and ChemiDoc XRS+ (Bio-Rad, CA, USA).

### Flow cytometry analysis

To measure proliferation, 2x10^4^ cells were seeded in 6-well plates, treated with 1 μM opioid, and counted by FACS each day for 7 days. For cell cycle analysis, cells were treated with 1 μM opioid for 24 h and subjected to cell cycle analysis by means of propidium iodide staining as previously reported (59). Analysis of apoptosis was performed using the Annexin V Apoptosis Detection Kit (eBioscience, MA, USA; 88-8006-74) according to the manufacturer’s protocol. To assess E-Cadherin cell surface expression, cells were stimulated for 1 h with 1 μM of the opioid following incubation with anti-E-Cadherin (dilution 1:1000, CST, 3199). Cell staining was recorded by a BD FACS Canto II flow cytometer and FACS Diva software (BD) as described (58).

### Reverse transcription PCR

Total RNA from control and opioid-treated cells was isolated using TRIzol (Peqlab, GER). cDNA was synthesized using iScript cDNA Synthesis Kit (Bio-Rad; 1708890). qPCR reactions were performed on a MyiQ2 cycler (Bio-Rad) with SsoAdvanced SYBR GreenSupermix (Bio-Rad). The following primers were used: *E-Cadherin* (NM_004360) Fw: 5’-*TCGCTTACACCATCCTCAGC*-3’ and Rev: 5’-*GGAAACTCTCTCGGTCCAGC*-3’, *SNAIL* (NM_005985) Fw: 5’-*TCCAGAGTTTACCTTCCAGCA*-3’ and Rev: 5’-*CTTTCCCACTGTCCTCATCTG*-3’, *SLUG* (NM_003068) Fw:5’-*ATATTCGGACCCACACATTACC*-3’ and Rev:5’-*GCTACACAGCAGCCAGATT*-3’, *STAT3* (NM_003150) Fw: 5’-*ACCAACAATCCCAAGAATGT*-3’ and Rev:5’-*CGATGCTCAGTCCTCGC*-3’, *TWIST* (NM_000474) Fw:5’-*GAGTCCGAGTCTTACGAGG*-3’ and Rev: 5’-*CCTCTACCAGGTCCTCCAGA*-3’. All gene expressions were normalized to *RPS18* (NM_022551) Fw: 5’-*ATTAGGGGTGTGGGCCGAAG*-3’ and Rev: 5’-*TGGCTAGGACCTGGCTGTAT*-3’.

### Cell migration

Migration of BCa cells was assessed by scratch assay. Therefore, cells were seeded onto 6-well plates at 5×10^4^ cells/well. Confluent monolayers were scratched with a sterile 1000 μl tip and washed two times with PBS. Cells were maintained in fresh DMEM medium and exposed to 1 μM opioid, naltrindole (selective DOR antagonist from Bachem) or ddH_2_O as vehicle control. Scratches were photographed at selected time points using an Olympus CK841 microscope (JP) and quantified with ImageJ software (60). For the transwell assay, 1×10^5^ cells were placed into an insert (Millicell, 8 μm pore size; Merck Millipore, MA, USA) with DMEM medium containing 1% FBS and 1 μM opioid, 3.3 nM ruxolitinib (Selleckchem, TX, USA) or ddH_2_O. Inserts were placed into a 24-well plate filled with DMEM medium containing 5% FBS as a chemoattractant. After incubation for 24 h at 37°C, cells that migrated through the membrane were fixed with 4% formaldehyde and 99% ice-cold methanol, and stained with DAPI (100 ng/ml). Insert membranes were imaged on an Olympus IX71 fluorescence microscope and migrated cells were quantified by ImageJ software.

### *In vivo* metastasis assay

*C.Cg-Rag2^tm1Fwa^ Il2rg^tm1Wji^* (*Rag2^-/-^γc^-/-^*) were backcrossed onto the BCa susceptible *Balb/C* background for more than six generations and maintained at the University of Veterinary Medicine Vienna under specific pathogen free (SPF) conditions. 1×10^6^ MDA-MB-231 were injected into the 4^th^ mammary fat pad of female *Rag2^-/-^γc^-/-^* mice or 1x10^4^ 4T1 cells were injected into the 4^th^ mammary fat pad of female wildtype mice. When tumors reached about 100 mm^3^ in volume, the primary tumors were resected. Mice were randomly divided into vehicle control (PBS) and opioid-treated groups (2 mg/kg opioid intraperitoneal injection; every 24 hours). Ten days after primary tumor removal, mice were sacrificed, lungs were isolated and the lung weight was assessed. Primary tumor volumes were calculated according to the formula: length × (diameter)^2^ × **π**/6.

### Histopathology

Lungs were fixed overnight in 4% Roti-Histofix (Carl Roth, GER), dehydrated, paraffin-embedded and cut in 4 μm sections. Sections were stained with Hematoxylin (Merck, GER) and Eosin G (Carl Roth). For immunohistochemical staining, heat-mediated antigen retrieval was performed in citrate buffer at pH 6.0 (Dako, CA, USA; S1699). Lung sections were then stained with antibodies against Cleaved Caspase-3 (dilution 1:200, CST; 9661), Ki-67 (dilution 1:1.000, eBioscience; 14.5698.80), Vimentin (dilution 1:200, CST, 5741) or CK8 (dilution 1:50, Leica microsystems, GER; NCL-L-CK8-TS1) using standard protocols. Images were taken using an Olympus IX71 microscope.

### Oncomine database

Gene expression datasets from human cancer (Bittner Breast, GSE2109; Miyake Breast, GSE32646; Okayama Lung, GSE31210; Cutcliff Kidney, GSE2712; Skotheim Testis, GSE1818; Quade Uterus, GSE764; Riker Melanoma, GSE7553; Ginos Head and Neck, DOI:10.1158/0008-5472.CAN-03-2144; Xu Melanoma, GSE7929; Bild Ovarian, GSE3149; Rickman Tongue, DOI:10.1038/onc.2008.251; Minn Breast, GSE2603; Desmedt Breast, GSE7390) were analyzed for the expression level of *OPRD1* by using Oncomine Compendium of Expression Array data (https://www.oncomine.org/resource/login.html) (61). Briefly, the *p* value for statistical significance was set at <0.05, while the fold change was set to 1.5 and the gene rank was defined as all.

### Statistics

Student’s t-test, one-way ANOVA and two-way ANOVA test were performed using GraphPad Prism^®^ Software version 5.04. Each experiment was performed in duplicate and repeated at least three times. The differences in mean values among groups were evaluated and expressed as the mean ± SD. *p* values less than 0.05 were considered statistically significant (^*^ p<0.05; ^**^ p<0.01; ^***^ p<0.001; ^****^ p<0.0001).

### Study approval

All experiments were approved by the institutional animal care committee and review board, and conform to Austrian law (BMBWF-68.205/0094-V/3b/2018).

## Author contributions

D.A.F and S.T. designed and supervised the study, S.T., H.A.N., V.M.K., and D.P.E. performed experiments. S.T., H.A.N. and D.P.E. analyzed data. F.A., D.A.F. and R.M. provided reagents and analytic tools. S.T., D.A.F and R.M. wrote the manuscript.

## Acknowledgments

D.A.F. was supported by the Austrian Science Fund (FWF), grant P27248-B28, R.M. and H.A.N. were supported by a private cancer metabolism grant donation from Liechtenstein and R.M. was further supported by the Austrian Science Fund (FWF), grants SFB F4707 and SFB-F06105, Austria and F.A. by FWF grants W1213, P25629, the priority program ACBN of the University of Salzburg and a smart specialization center grant by the County of Salzburg.

We thank Veronika Sexl and Sabine Mascho-Machler for scientific input. For technical support we thank Michaela Prchal-Murphy, Stephanie Kaspar, Safia Zahma and Sabine Fajmann, plus the animal facility team.

